# Nerve injury triggers nociceptive hypersensitivity with interhemispheric divergence in haplodeficient GAD67-GFP mice

**DOI:** 10.64898/2026.05.17.725734

**Authors:** Johannes Spahn, Clara A. Simacek, Lisa Hahnefeld, Luisa Franck, Marc-Philipp Weyer, Chloe Hall, Robert Gurke, Thomas Mittmann, Irmgard Tegeder

## Abstract

Nerve injury causes an imbalance of glutamatergic excitation over GABAergic inhibition, contributing thereby to lasting neuropathic pain. Transgenic GAD67-GFP knock-in reporter mice were developed to visualize GABAergic interneurons. The knock-in into glutamate decarboxylase (GAD67) causes haploinsufficiency that manifest in low GABA levels. In this model, we studied if diminished GABA exacerbates neuropathic pain after nerve injury. Adolescent male and female GAD67-GFP knock-in mice were subjected to Spared Sciatic Nerve Injury (SNI). At baseline, nociception and thermal preferences were equal but after SNI, GAD67-GFP mice developed thermal allodynia which was not detected in wildtype littermates. At the electrophysiology level, SNI caused a partial decrease in the excitability in layer 2/3 pyramidal neurons in the projection-hemisphere in wildtype mice. This effect was exacerbated in GAD67-GFP, affecting both sides, and was accompanied with imbalance of field-potential (FP) amplitudes between projection and non-projection hemisphere, which did not occur in wildtype mice. The results suggest that GABA deficiency can be compensated to protect from hyperexcitability at baseline, but it cannot be further upscaled, ultimately leading to network hyperactivity after injury. Metabolomic studies confirmed the moderate loss of GABA in ipsi- and contralateral cortex and lumbar spinal cord of GAD67-GFP mice and failure to raise GABA in the ipsilateral dorsal horn after injury. Carnosine, cystathionine, and glutathione, three important anti-oxidative metabolites, were co-reduced with GABA suggesting that GABAergic activity and anti-oxidative capacity are interconnected and failure of this axis contributes to neuropathic “pain”.

## Introduction

Neuropathic pain is caused by diverse insults to the peripheral or central nervous system including trauma, toxic damage, inflammation, infection, autoimmunity or metabolic disorder [1]. It may affect single nerves or manifests as polyneuropathy. Therapeutic options have increased over the last decade but are still limited and insufficient in about one third of patients who seriously suffer from persistent intricate pain, rather than from the underlying disease or injury [2, 3]. It is increasingly recognized that the endogenous mechanisms of pain control involve opioid-[4, 5], and cannabinoid-dependent networks [6], inhibitory GABA- and glycine-ergic [7, 8], as well as adrenergic [9] and peptidergic [10] circuits, which determine the efficacy of pharmacologic and non-pharmacologic approaches.

Inhibitory interneurons play a crucial role in the development of neuropathic pain [11-13]. It is believed that deficiency of gamma aminobutyric acid (GABA) in the spinal cord, somatosensory cortex and pain-associated regions of the “pain-matrix” may increase the risk for the development of neuropathic pain [14-16], because GABA is one of the major inhibitory neurotransmitters in nociceptive and pain-associated circuits [17]. GABA is the main inhibitory neurotransmitter in the mature CNS [18-20], and the gradual maturation of the GABAergic system determines the timing of the critical periods for sensory system development and synapse maturation [21-24]. A reduced level of GABA can strongly impact brain functions both during development and in adulthood [25-27]. Under healthy conditions the brain balances the strengths of glutamatergic excitation (E) versus GABAergic inhibition (I), establishing an appropriate E/I balance, which can be seriously disrupted throughout life by various insults [28-30] or aging [31]. In humans, such imbalances have been associated with neuropsychiatric diseases like autism-spectrum disorders (ASD) and schizophrenia, epilepsy [32-36] and traumatic brain injury (TBI) and post TBI epilepsy [30, 37-39]. However, in the early phase of TBI, compensatory mechanisms are able to rebalance the E/I ratio in the cortex [30] suggesting that similar mechanisms may occur upon peripheral nerve injury that might be enhanced or supported therapeutically to combat neuropathic pain.

GAD67-GFP knock-in mice are a well-established mouse model to study GABAergic interneurons in living tissues [40]. This mouse line significantly contributed to a better understanding of the role of GABAergic interneurons in normal brain development and also under pathological conditions like autism spectrum disorder, schizophrenia or TBI [30, 41, 42]. However, GFP labeling causes the loss of one functional copy of the GAD1 gene, encoding the GAD67 glutamate decarboxylase enzyme, the main producer of GABA in the developing CNS [43]. As a consequence, the lack of one functional copy of the GAD1 gene leads to a reduced functional GAD67 enzyme activity and in turn a decreased GABA concentration in the CNS [40]. Indeed, electrophysiological or behavioral studies in GAD67-GFP knockin mice demonstrated changes in E/I balances [44, 45] and social behavior [46] or attention deficit disorder-like behavior [47], but interestingly, the mice did not develop epileptic seizures [40], which indicates that compensatory mechanisms kick in to prevent chronic hyperexcitability and epileptogenesis.

Our previous studies in GAD67-GFP mice have shown that GABA levels are indeed moderately reduced in the developing cortex of these mice, and whole-cell patch clamp recordings revealed that the frequency of miniature inhibitory postsynaptic currents (mIPSCs) was reduced in layer 2/3 pyramidal neurons at P14 and at P21 without any change in amplitude or kinetics [48]. Overall, the E/I-ratio was shifted towards excitation. However, multielectrode-recordings (MEA) from acute slices revealed a decreased spontaneous neuronal network activity compared to wildtype (WT) littermates, pointing to a compensatory mechanism that prevented seizure-like hyperexcitability and depended on a tonic activation of GABA-B receptors [48]. This was possible because GABA transporter 3 (GAT-3, Gene Slc6a11) acted in a reverse mode, leading to GABA release instead of uptake [48]. This important endogenous mechanism could determine the outcome of nerve injury associated neuropathic pain [35, 49]. Hence, the operation mode of GAT3 may determine if pain resolves, or if this mechanism might be sustained therapeutically to fortify the pain defense. It has been reported previously that intrathecal delivery of a GAT3 inhibitor reduces acute nociception in rats [50] but its inhibition is unlikely to provide a lasting therapeutic benefit.

To further investigate the role of GABAergic inhibition in adolescent brain and axonal injuries we used hemizygous GAD67-GFP mice and subjected them to an axonal injury of the sciatic nerve by transection of one of its branches (tibial). We hypothesized that GAD67-GFP mice would have “normal” nociceptive function and sensitivity at baseline but develop a stronger injury phenotype. If the hypothesis was true, sustaining GAT3 reverse transport might be useful to attenuate neuropathic pain.

### Methods

### Mice

Adolescent transgenic heterozygous GAD67-GFP positive female and male knockin mice (n = 7/9 male/female) and their GAD67-GFP negative wildtype (WT) littermates (n = 8 f/m) were used. Mice were three weeks old (24 x 22 days, 6 x 24 days, 2 x 27 days) at baseline nociception and were subjected to SNI surgery in the afternoon or the following morning after baseline thermal gradient ring (TGR) observation. Post SNI TGR was done 7 days after injury (7 dpi). Body weights were monitored before and after surgery, at 1 and 7 dpi and before euthanization.

The mouse line was originally generated by Tamamaki et al. [40]. For the present experiments we crossed knockin with wildtype C57BL6N mice. At birth, brains were optically inspected for presence or absence of GFP-positive signals by use of a fluorescent lamp, and the mice were labeled accordingly by toe clippings. For genotyping, final ear punch tissue was used in a Transnetyx assay. The genotyping result agreed 100% with the fluorescent lamp genotyping. Animals were kept under a standard 12h day/night rhythm at a room temperature of 22-24°C with an ad libitum supply of food and water. The experiments were designed to restrict the number of animals that were used in this study to the necessary minimum, and all experiments were performed in accordance with German and European laws of animal welfare in science. The experiments were approved by the Regierungspräsidium Darmstadt under the approval number FK1110.

### SNI (spared nerve injury)

Surgery was performed in 3-4 weeks old adolescent mice under 1.5-2 % isoflurane anesthesia with local lidocaine anesthesia of the skin as described [51, 52]. To induce a one-branch spared nerve injury of the sciatic nerve (SNI), the tibial branch of the left sciatic nerve was constricted with 6/0 silk suture and distally transected. The sural and the peroneal branches were spared. One-branch SNI was used instead of the more common 2-branch SNI because in adolescent test mice, cutting of both, the tibial and peroneal branches, caused a serious impairment of walking, which did not occur with tibial-only injury. One-branch models have been shown to be preferable for mechanistic studies [53, 54]. The tibial nerve stump and the correct positioning of the suture few millimeters proximal of the stump end was confirmed by necropsy and histology of the sciatic nerve.

SNI including one-branch SNI [54] causes hyper-excitability in the ipsilateral dorsal horn of the spinal cord, i.e. at the first synapse of the injured sensory nerve. In wildtype mice, GABA is increased in the ipsilateral dorsal horn after sciatic nerve injury which is an endogenous mechanism of pain defense [55]. We hypothesized that the injury evoked GABA raise was not effective in GAD67-GFP mice, which may lead to exacerbation of hyperexcitability and nociceptive hypersensitivity.

### Assessment of thermal sensitivity using a Thermal Gradient Ring (TGR)

A thermal gradient ring (TGR, Ugo Basile https://ugobasile.com/products/categories/pain-and-inflammation/new-thermal-gradient-ring-tgr-zimmermann-s-method-2-0) was used to assess the temperature preferences and the exploration of the ring platform that consists of a circular runway allowing free choice of the comfort zone [56, 57]. The dimensions of inner and outer ring diameters are 45 cm and 57 cm. The inner walls consist of plexiglass and the outer walls of aluminum. Both are 12 cm high and build a 6 cm wide circular running arena. The aluminum surface provides a temperature gradient that is controlled with two Peltier elements and constantly measured with infrared cameras. The arena is divided into mirror-image semicircles of 12 temperature zones, so that duplicate readouts are provided for each zone. The temperature zones were set to range from 14°C-40°C. During measurements, the running track was illuminated, and the mouse behavior was videotaped with a regular CCD camera, mounted above the mid-point of the ring. The time spent in zones and temperature preferences, walking distances and speed, and zone entries and rotations were analyzed with the TGR ANY-Maze video tracking software (Stoelting). In addition, rearing and grooming were monitored in the first 15 min by pressing defined keys. TGR monitoring was done before SNI surgery and 7 dpi, each with 60 min observation.

### Mice tissue collection: brain and plasma

Mice were euthanized with slowly rising carbon dioxide and blood withdrawal by cardiac puncture, whereby the blood was collected into K3 EDTA tubes, centrifugated at 1300 *g* for 5 min, and plasma was transferred to a fresh tube and snap frozen on dry ice or in liquid nitrogen. The ipsilateral and contralateral brain and lumbar spinal cord were dissected for lipidomic and metabolomic analyses. Cerebellum and olfactory bulb were removed, and the brain was cut sagittal. Left and right halves were weighed with precision scales and snap frozen on dry ice. Subsequently, the lumbar spinal cord was dissected and processed accordingly. Finally, the sciatic nerves were dissected, washed in 1x PBS and then fixated in 4% paraformaldehyde (PFA) for histology. Samples were stored at -80 °C until analysis.

### Electrophysiology

After completion of SNI and the behavioral experiments, a subgroup of the animals of both genotypes at P34 – P40 were examined via multi-electrode arrays and whole-cell patch-clamp recordings in acute brain slices.

#### Slice Preparation

The slice preparation protocol was based on a protocol provided by Ting et al. [58]. Mice were terminally anaesthetized with 4% isoflurane and decapitated. After careful removal, the brains were transferred into ice cold and oxygenated (95% O2, 5% CO2) artificial cerebrospinal fluid (aCSF), enriched with the protective reagent N-methyl-D-glucamine (NMDG), containing (in mM): NMDG, 92; NaH2PO4, 1.25; NaHCO3, 30; HEPES, 20; Glucose, 25; thiourea, 2; Na-ascorbate, 5; Na-pyruvate, 3; CaCl2, 0.5; MgSO4, 10; KCl, 2.5; pH adjusted to 7.3 – 7.4 (all substances obtained from Carl Roth, Karlsruhe, Germany). Next, 300 µm thick coronal brain slices were cut using a vibratome and hemispheres were bisected using fine needles. Slices of the ipsi- and contralateral hemisphere containing the somatosensory cortex were incubated in separate small containers for 25 min at 37°C in NMDG-aCSF. During this period, a Na^+^-based solution (2.32 g NaCl dissolved in 20 ml NMDG-aCSF) was added every 5 minutes within 25 minutes total (Volumes: 333 µl, 333 µl, 667 µl, 1333 µl, 2667 µl). Brain slices were then transferred into standard aCFS (containing in mM: NaCl, 125; KCl, 2.5; MgSO4 × 7 H2O, 1; CaCl2 × H2O, 2; NaH2PO4 × H2O, 1.25; NaHCO3, 25 and d-glucose, 25; pH: 7.4, Carl Roth, Germany) for a further incubation for at least 45 min at room temperature.

#### Whole-Cell Patch-Clamp Recordings

For patch-clamp recordings, brain slices were placed on a submerged recording chamber and constantly perfused with oxygenated aCSF at 37°C. The chamber was mounted on an upright microscope (Olympus BX51WI, Olympus, Tokyo, Japan) and the primary somatosensory cortex was identified using an objective lens of lower magnification (2.5x, Olympus, Tokyo). Pyramidal neurons were visually identified by their morphology using an objective lens of higher magnification (40x, Olympus). All electrophysiological experiments were conducted with an Axopatch-200B amplifier and Clampex 11.2 software (Molecular Devices, San José, CA, USA). Borosilicate glass pipettes (GB 150F-8P; Science Products, Frankfurt, Germany) with resistances between 7 – 10 MΩ were produced with a DMZ Zeitz-Puller (Planegg, Germany). Recordings of passive and active membrane properties were conducted using intracellular solution I) containing (in mM): K-gluconate, 140; KCl, 8; MgCl_2_ × 6 H_2_O, 2; Na_2_ATP, 4; Na_2_GTP hydrate, 0.3; Na_2_Phosphocreatin, 10; HEPES potassium salt, 10. For experiments of excitatory and inhibitory postsynaptic currents (sEPSCs and sIPSCs, respectively) intracellular solution II) was used containing: Cs-gluconate, 125; CsCl, 5; EGTA, 10; MgCl2, 2; Na2ATP, 2; Na2GTP, 0.4; HEPES, 10. The pH values were adjusted to 7.3 using KOH for solution I) and CsOH for solution II). The serial resistance (R_s_) and membrane resistance (R_m_) were measured before and after every experiment, and cells were excluded from further analysis, when R_s_ exceeded 20 MΩ or was changed >20% between the recordings. Recordings were filtered at 2 kHz and digitized at 50 kHz using a Digidata-1400 system and Clampex 11.1 software (Molecular Devices, San José, CA, USA).

#### Passive and active membrane properties

The resting membrane potential was recorded right after cell membrane rupture, and neurons were subsequently clamped at a potential of -70 mV. Membrane resistance (R_m_) and the membrane capacitance (C_m_) were determined by first injecting a hyperpolarizing current of -10 pA for 1000 ms (20 repetitions), followed by fitting a mono-exponential curve. Active membrane properties were obtained from the first action potential at rheobase, elicited by the application of a current step protocol (from -100 pA to +500 pA; step size of 10 pA; step duration of 1000 ms). Active membrane properties such as the action potential (AP) firing threshold, AP-amplitude, as well as the AP-firing frequency were analyzed at the first AP at rheobase using Clampfit11 software (Molecular Devices).

### Excitatory and inhibitory postsynaptic currents

Recordings of spontaneous excitatory and inhibitory postsynaptic currents (sEPSCs and sIPSCs, respectively) were performed using the Cs-based intracellular solution II) to block potassium conductance. All recordings were conducted in voltage-clamp mode and after cell membrane rupture, cells were held at -70 mV. For measuring sEPSCs, cells were clamped at the Nernst-reversal potential of GABA_A_-receptors (-60 mV). To isolate AMPA-receptor-mediated currents, 2-amino-5-phosphonovalerinans acid (DAP-5, 25 μM) was added to the aCSF. sIPSCs were measured at a holding potential of +10 mV, the Nernst-reversal potential of AMPA-receptor-mediated currents, to isolate GABA_A_-receptor-mediated currents. No serial resistance compensation was applied and all recordings of sEPSCs and sIPSCs lasted for 5 min. Analysis was conducted using MiniAnalysis software (Synaptosoft, Fort Lee, NJ, USA) and files were blinded during analysis.

### Multi-electrode-array (MEA) recordings

Spontaneous cortical network activity was recorded in the acute slices by a MEA system consisting of two recording chambers (MEA2100 System, Multi-Channel Systems MCS GmbH, Germany). Each MEA chip had 60 electrodes (60MEA200/30iR) with an electrode-diameter of 30 μm and an inter-electrode distance of 200 μm. Cortical slices were placed on the MEA aligning the outer cortical border along the first row of electrodes. The electrode rows with the number 2 and 3 corresponded to the cortical layer 2/3 and were used for further analysis. Each MEA chip was used for multiple recordings of several brain slices in randomized alternating order of the mice and without knowledge of the genotype. Genotyping was done post-MEA with ear punches obtained during SNI surgery. Slices were carefully placed and secured on the MEA chip using a platinum grid and constantly perfused with oxygenated recording aCSF at a flow rate of 1.5 ml/min using a centrifugal pump (Gilson international, Berlin, Germany). The recording chamber containing ACSF was heated to 32^°^C. The slices equilibrated on the chip for 30 min before starting the electrophysiological recordings. Raw data were recorded using a sampling frequency of 50 kHz and filtered using a Butterworth high-pass second order filter with a 200 Hz cut-off. Input-output curves were generated from increasing electrical stimulations in steps of 500 mV, from 500 mV to 5000 mV, at time intervals of 40 s and a total of three rounds of stimulation per slice. The peak value of each response per input was analyzed using MultiChannel Analyser 2.2 software and the mean was taken across the repeats.

### Untargeted lipidomic and metabolomic analyses

Mouse tissue samples were homogenized by adding ethanol:water (1:3, v/v, tissue concentration 0.02 µg/ml) using a Precellys 24-dual tissue homogenizer coupled with a Cryolys cooling module (both Bertin Technologies, Montigny-le-Bretonneux, France) with 10 zirconium dioxide grinding balls (3*20s at 6500 g with 60s breaks), operated at < 6 °C. Subsequently, 20 µl (cortex) or 40 µl (spinal cord) of the homogenate containing 1 mg of tissue were extracted using a liquid-liquid-extraction method.

Lipidomic and metabolomic analysis were conducted applying the same procedure as previously described [59]. Further protocol details for extraction and analyses of different matrices (brain, spinal cord) are described in the supplementary methods (Excel file).

Briefly, a methyl-tert-butyl-ether (MTBE) and methanol-based liquid-liquid extraction was used to allow for the simultaneous analysis of polar metabolites and lipids from the same sample.

For chromatographic separation of lipids, a Zorbax RRHD Eclipse Plus C8 1.8 µm 50 x 2.1 mm ID column (Agilent, Waldbronn, Germany) with a pre-column of the same type was used. The mobile phases were (A) 0.1% formic acid and 10 mM ammonium formate and (B) 0.1% formic acid in acetonitrile:isopropanol (2:3, v/v). For metabolomic analysis, polar metabolites were separated on a SeQuant ZIC-HILIC, 3.5 µm, 100 mm × 2.1 mm I.D. column coupled to a guard column with the same chemistry (both Merck, Darmstadt, Germany) and a KrudKatcher inline filter (Phenomenex, Aschaffenburg, Germany). Using 0.1 % formic acid in water (solvent A) and 0.1 % formic acid in acetonitrile (solvent B), binary gradient elution was performed with a run time of 25 min.

Lipids and polar metabolites were analyzed on an Orbitrap Exploris 480 with a Vanquish horizon UHPLC system (both Thermo Fisher Scientific, Dreieich, Germany). Data was acquired using Thermo Scientific XCalibur v4.4 (RRID:SCR_014593) and relative quantification was performed in Thermo Scientific TraceFinder 5.1 (RRID:SCR_023045). Full scan spectra were acquired from 180-1500 *m/z* (lipidomics), 70-700 *m/z* (metabolomics positive ion mode) or 59-590 *m/z* (metabolomics negative ion mode) at 120,000 mass resolution each for 0.6 sec, and data dependent MS/MS spectra at 15,000 mass resolution in between.

For all lipidomic and metabolomic analyses, the area under the curve (AUC) was used for quantification. AUCs were normalized using a median-based probabilistic quotient normalization (PQN) method to adjust for technical runtime and matrix differences of different source tissues (brain, spinal cord). AUCs were transformed to the square root of the AUC (sqrt AUC or sqrt AUC/IS) for data analysis to adjust skewed distributions. For multivariate analyses, sqrt AUCs were further scaled to have a common mean and variance of 1 (autoscaling in MetaboAnalyst).

### Data analyses and statistical procedures

Group data are presented as mean ± SD, mean ± sem for behavioral data, or median ± IQR as specified in the respective figure legends. Data were exported from acquisition or recording software to Excel and further analyzed with SPSS 29 (RRID:SCR_016479), GraphPad Prism 9 or 10 (RRID:SCR_002798), Origin Pro 2025 (RRID:SCR_014212), and MetaboAnalyst 5.0 (RRID:SCR_016723) (https://www.metaboanalyst.ca) [60]. Area under the curve (AUC) data of lipidomic and metabolomic analyses were transformed to square root (sqrt) values to adjust skewed distributions and were PQN-normalized to adjust for differences in extraction efficiency with different tissue matrices. For testing the null-hypothesis that groups were identical, two groups were compared with 2-sided, unpaired Student’s t-tests. Electrophysiology, lipidomic and metabolomic data were submitted to one-way or two-way analysis of variance (ANOVA) using e.g., the factors “feature” (e.g. lipid, metabolite) and ‘group’ (e.g. genotype) and time point (before/after SNI). In case of significant differences, groups were mutually compared using post hoc t-tests according to Tukey (E-phys), Šidák (behavior) or false discovery rate (FDR) (omic data). The meaning of asterisks in figures is explained in legends. The graphs only show results of P-values < 0.05 (asterisks) or < 0.1 (exact P-values). Partial Least Square discrimination analysis (PLS-DA) was used to assess the prediction of group membership and classification according to the variable importance. For multivariate analyses, data were normalized to have a common average and variance of 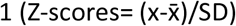 “autoscale” in MetaboAnalyst).

## Results

### General health and GABA levels in GAD67-GFP mice

Hemizygous GAD67-GFP mice were previously shown to generate reduced concentrations of cortical GABA at P14 [40], which was confirmed in our mouse colony previously [48]. In the present study GABA levels were measured in ipsi- and contralateral brain and spinal cord by UHPLC-MS/MS metabolome studies to assess the importance of GABA homeostasis in the young adult CNS in the metabolic context of the brain and spinal cord. The expected reduction of GABA was confirmed (see below). Although our previous electrophysiological study *on ex* vivo slices in juvenile mice showed a reduced frequency of miniature inhibitory postsynaptic currents (mIPSCs) indicating a weaker phasic GABAergic drive [48], the adolescent GAD67-GFP mice of the present study did not show symptoms of hyperexcitability at the behavioral and general health levels. Body weights were equal to wildtype littermates throughout our experiments before and after SNI surgery, both in male and female mice (Figure 1).

**Figure 1.**
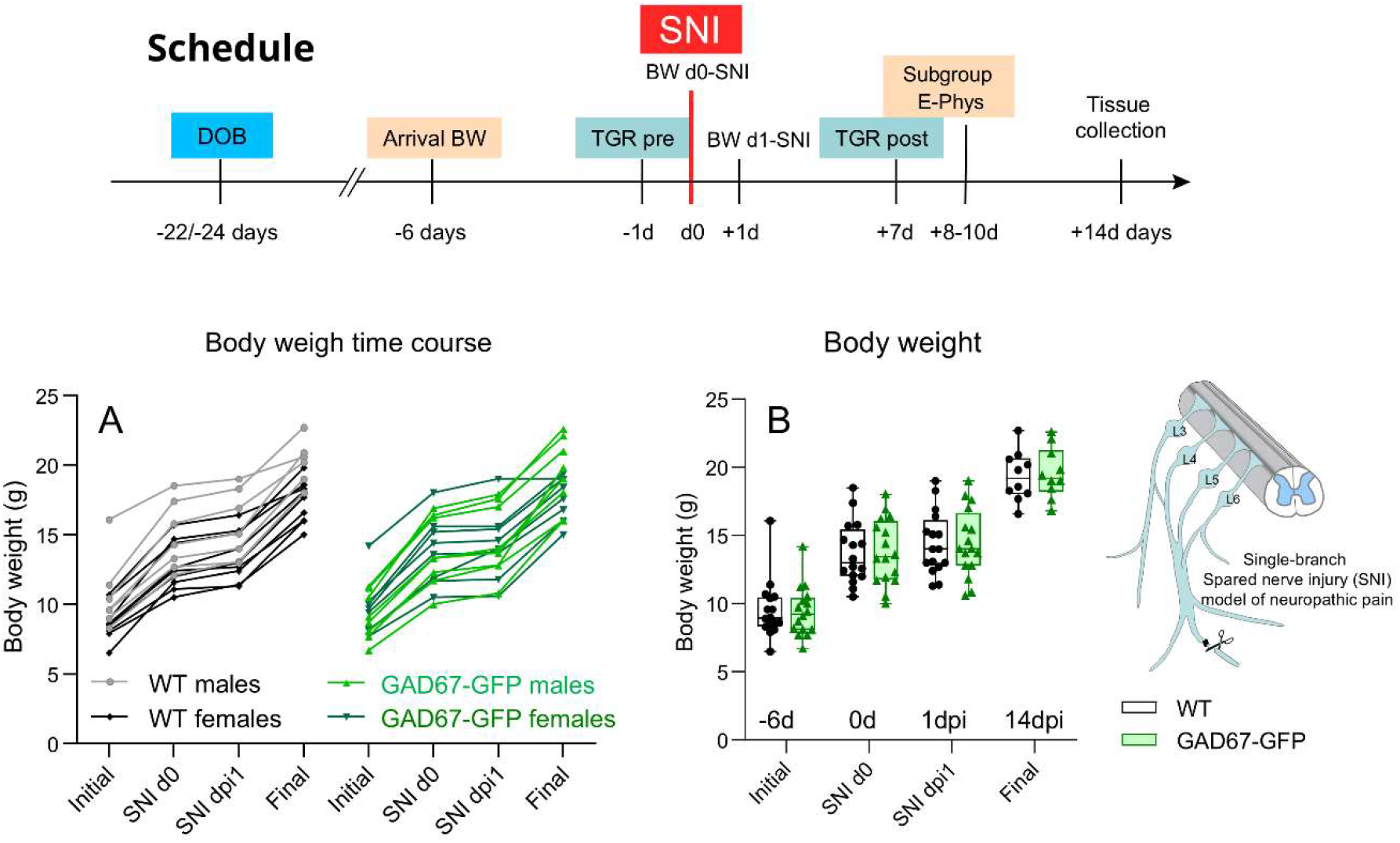
Schedule and body weights before and after Spared Nerve Injury of the sciatic nerve (SN) in adolescent mice **A:** Individual body weight time courses in male and female GAD67-GFP and wildtype control mice. The ages at the initial time point 6 days before surgery were 22d (n = 12/12), 24d (n=3/3) and 27d (n = 1/1) equally distributed between genotypes. **B:** Box/scatter plots of body weights per genotype. Body weights increased after SNI. Surgery did not cause weight loss. Data were compared with two-way ANOVA for the within subject factor “time point” (i.e. before/after SNI) and the between subject factor “genotype” and subsequent posthoc analysis for genotype. There were no statistical differences between groups.

### Sensorimotor behavior in the Thermal Gradient Ring

A Thermal Gradient Ring was used to investigate cold and heat sensitivity and thermal allodynia that develops in mice after SNI surgery [52, 61]. It is in an unbiased video-based test and showed normal baseline behavior (sensory and motor) in GAD67-GFP mice but thermal allodynia 7 days after SNI (7 dpi). Exemplary short video-traces are included as supplementary files. To assess the exploratory activity, rearing behavior was recorded in the first quarter of the test (Figure 2A). Most of the mice showed an increase of the rearing time in the second test at 7 dpi as compared to the baseline test which indicates a recognition of the familiarized TGR platform and maybe a manifestation of post-SNI restlessness. Accordingly, the total distances travelled in the TGR test 7 dpi was shorter than at baseline (Figure 2B), but there was no difference between genotypes neither for rearing nor travel distance at baseline or after SNI (Figure 2A, B). The activity in the TGR followed a typical time-dependent pattern in both genotypes with highest activity and exploration in the first quarter, followed by settling down and resting in the preferred zone in quarter-2 and quarter-3 and a rerise of the activity in the last quarter. The timeline was very similar in both genotypes (Figure 2B).

**Figure 2.**
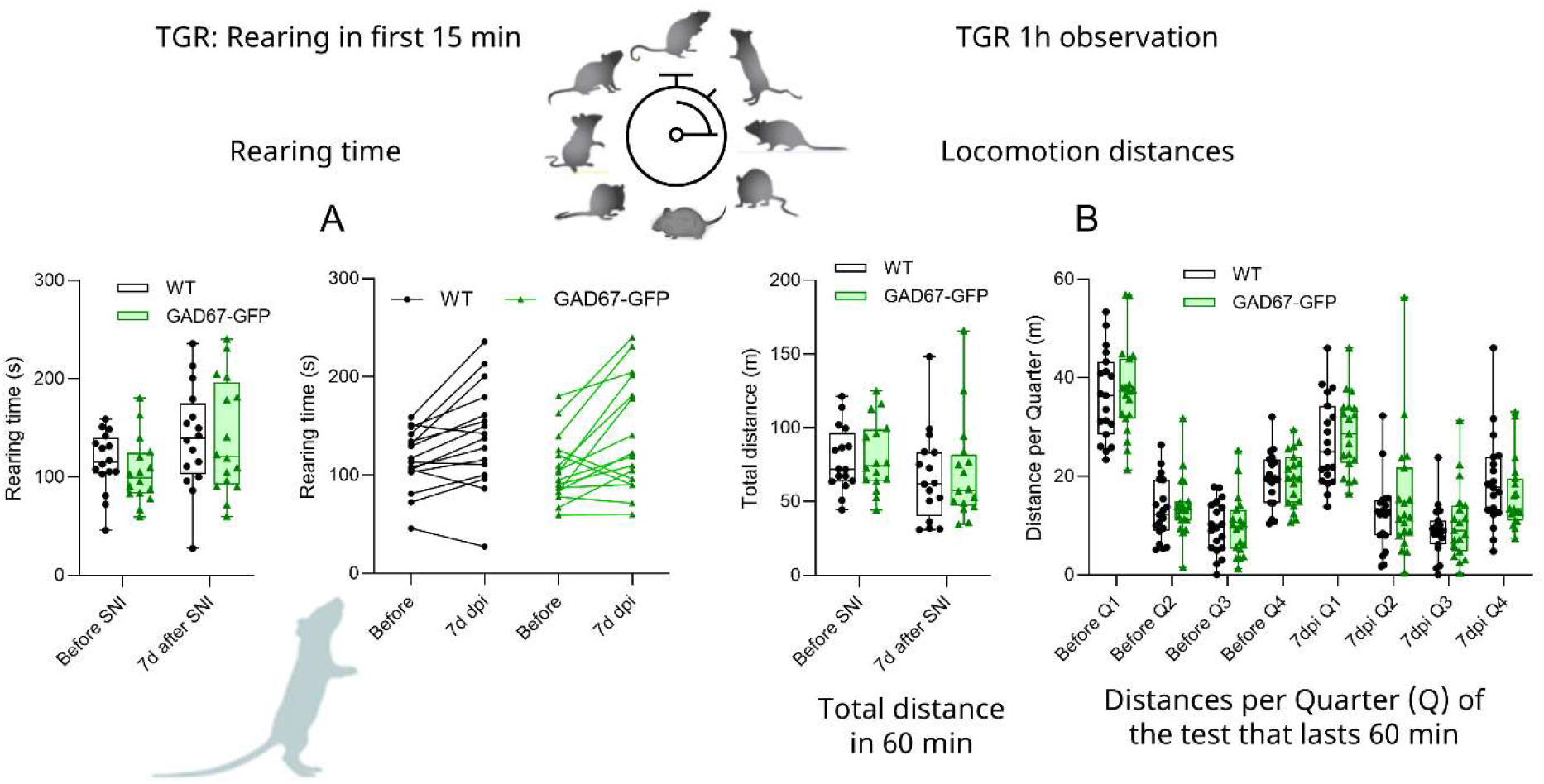
Motor behavior in the Thermal Gradient Ring in GAD67-GFP and wildtype mice before and after SNI **A:** Box/scatter and paired line graphs of the rearing time in the first quarter of the TGR test before and after SNI in GAD67-GFP and wildtype mice. The rearing behavior in the post-SNI TGR test increased as compared to the pre-surgery TGR-test in both genotypes indicating recognition of the circular runway and platform in the second test that gives rise to a decrease of horizontal but increase of vertical exploration, manifesting in an increase of the rearing time. **B:** Travel distances in the circular TGR platform gangway before and after SNI. While vertical exploration increases (A), horizontal exploration decreases in the second trial which was 7 days after injury resulting in a decrease of travel distances in both genotypes. In each trial, motor activity was highest in the first quarter, followed by resting in the comfort zone and re-rise in the last quarter. Data were compared with two-way ANOVA for the within subject factor “time point” (i.e. before/after SNI) and the between subject factor “genotype” and subsequent posthoc analysis for genotype. The boxes show the interquartile range, the line is the median, the whiskers show minimum to maximum and the scatters are individual mice. There were no statistically significant differences between genotypes.

The preference temperature ranged from 29.5-31.5 °C (interquartile range) in both genotypes at baseline (Figure 3A) and the average remained stable in wildtype mice after SNI whereas the preference temperature dropped by 2°C in GAD67-GFP mice (IQR 26.5-29.5) at 7 dpi after SNI indicating thermal hyperalgesia or allodynia. In three mice, the preference temperature was below 25 °C, substantially cooler than the skin temperature (about 29-30.5 °C), suggesting allodynia in these mice. Five (3m, 2f) of the WT mice showed an increase of the preference temperature after SNI suggesting that sensory loss or numbness still prevailed in these mice (Figure 3A). The initial sensory loss after SNI is normally replaced or overshadowed by hypersensitivity and not detectable with paw-withdrawal based tests.

**Figure 3.**
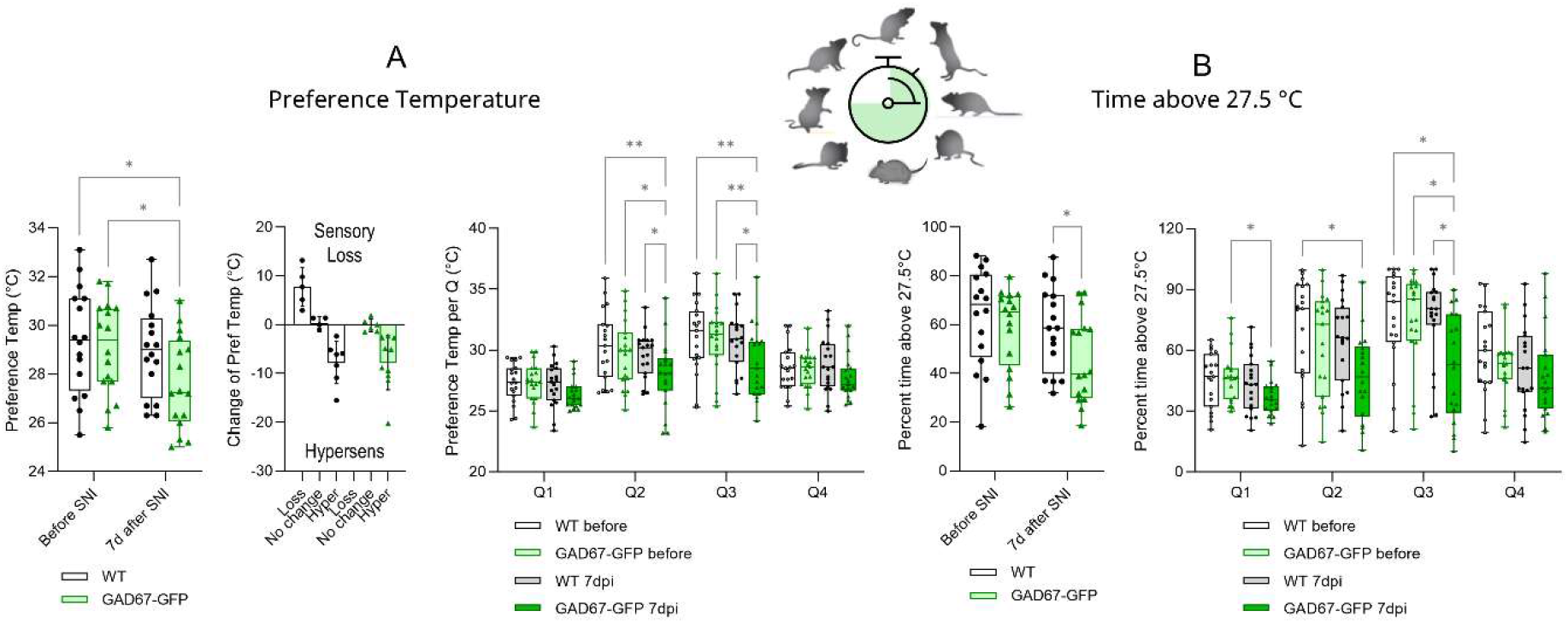
Preference temperatures before and after SNI in GAD67-GFP and wildtype mice **A, B:** Before SNI, preference temperatures and the times spent in zones above 27.5 degrees Celsius were equal in both genotypes. After SNI, average preference temperatures and times > 27.5°C remained stable in wildtype mice but dropped significantly in GAD67-GFP who avoided pleasant warm temperatures after SNI, mainly during the first 3 quarters of the TGR test. The behavior indicates thermal hypersensitivity and allodynia. Data were compared with two-way ANOVA for the within subject factor “time point” (i.e. before/after SNI) and the between subject factor “genotype” and subsequent posthoc analysis using an adjustment of alpha according to Tukey. The boxes show the interquartile range, the line is the median, the whiskers show minimum to maximum and the scatters are individual mice. Asterisks indicate statistically significant differences between genotypes. *P<0.05; **P<0.01.

The differences between genotypes were also evident in the “time spent above 27.5°C” (Figure 3B) and were stronger in the first 3 quarters of the test because the increase of the activity in the last quarter partially masks the distinction of the preferred zone (Figure 3).

To further assess the distribution of the preference temperatures the times spent in zones was plotted versus the mid-temperature of the zone for each quarter of the test (Figure 4). The distributions are characteristic for each quarter but almost equal between the genotypes at baseline (Figure 4A). However, after SNI the curves are flattened in GAD67-GFP mice and shifted to lower temperatures again clearly indicating thermal hypersensitivity (Figure 4B).

**Figure 4.**
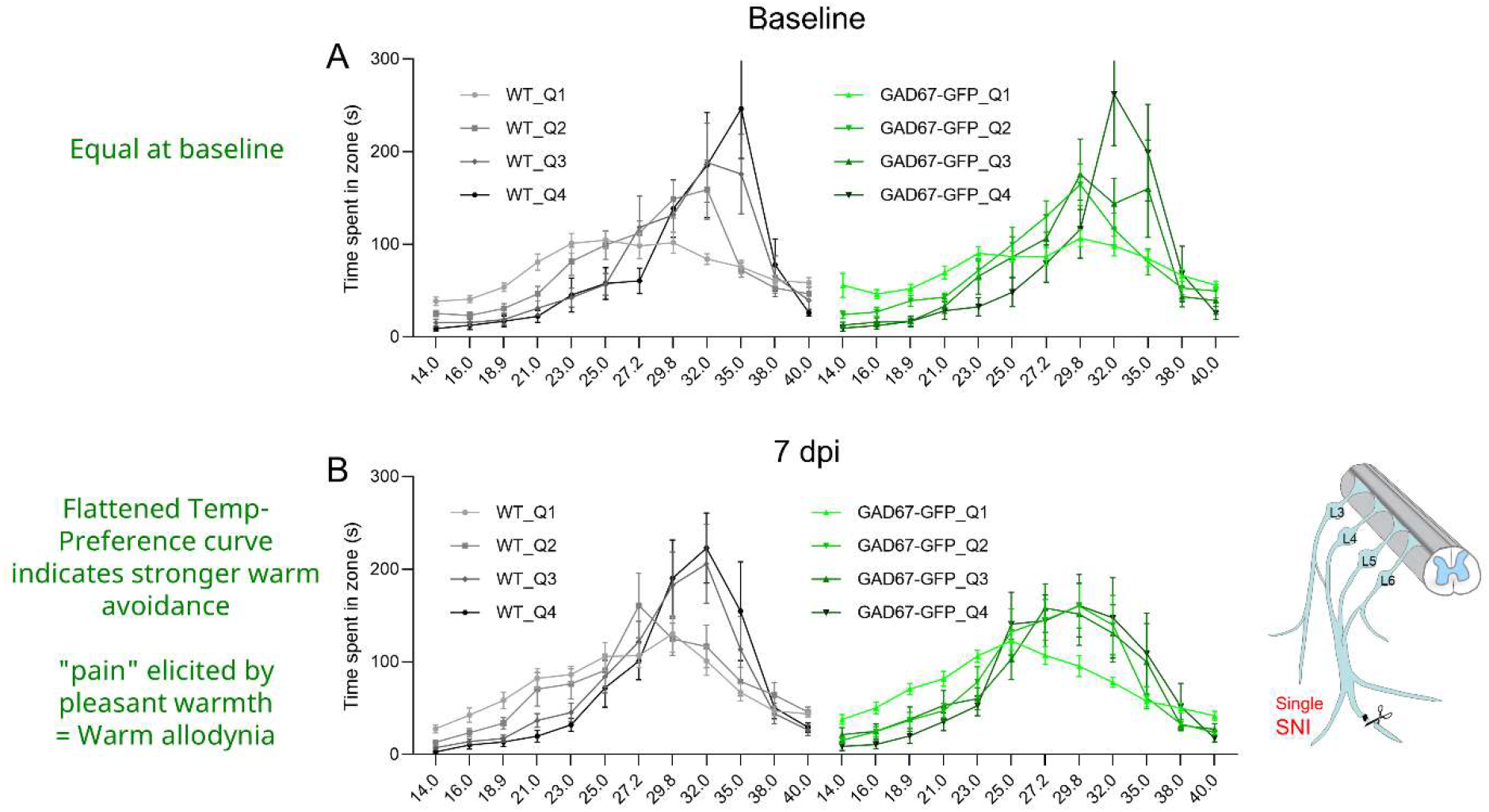
Temperature preference curves before and after SNI in GAD67-GFP and wildtype mice. **A:** Times spent in the temperature zones (Y-axis) in dependence of the temperature of the zone (x-axis) for each quarter of the TGR test before SNI. The curves get steeper with each quarter showing development of the preference of a warm comfort zone, equal in both genotypes. **B:** In analogy to A the curves show the times spent in temperature zones after SNI. While the preferences of wildtype mice after SNI are about equal to the baseline test, GAD67-GFP mice show flattened preference curves showing that they did not develop a preference for warm but rather avoided pleasant warm temperatures, again indicating post-SNI thermal allodynia. Statistics in Figure 3.

### Reduced activity of pyramidal neurons in GAD67-GFP mice

We hypothesized that thermal nociceptive hypersensitivity of GAD67-GFP mice would be associated with an increase of the neuronal activity in the somatosensory cortex reflecting the pain-associated glutamatergic neuronal hyperactivity, exacerbated by the lack of GABA. To explore this, we assessed putative injury-induced changes in functional properties of cortical pyramidal neurons in wildtype and GAD67-GFP mice. The mean resting membrane potential and AP threshold of pyramidal neurons of both hemispheres were neither affected by injury nor genotype (Figure 5A, B). Interestingly, the membrane resistance was reduced in GAD67-GFP mice, and the membrane capacitance remained constant in both genotypes (Figure 5C and D). This indicates a partial reduction of the intrinsic excitability in GAD67-GFP mice while the speed of signal integration remains constant. The reduced excitability was also reflected in a decreased firing frequency and action potential amplitude (Figure 5E-H). In addition, the mean membrane resistance and firing frequency were reduced in the contralateral hemisphere (right hemisphere) of WT mice (Figure 5 C, H). This part of the cortex receives direct input from the injured left nerve, whereas the left ipsilateral wildtype hemisphere receives no direct input and is therefore considered as the “healthy” side (Figure 5A, B). We infer that the injury-induced decrease in membrane resistance and AP firing frequency is an adaptive injury-evoked mechanism possibly contributed by loss of the sensory input from the innervation area of the injured branch. Further, our data suggests that the GAD67-GFP mice lack this adaptive capability, since their membrane resistance and firing frequency were already reduced under healthy conditions, and no further upscale was possible.

**Figure 5.**
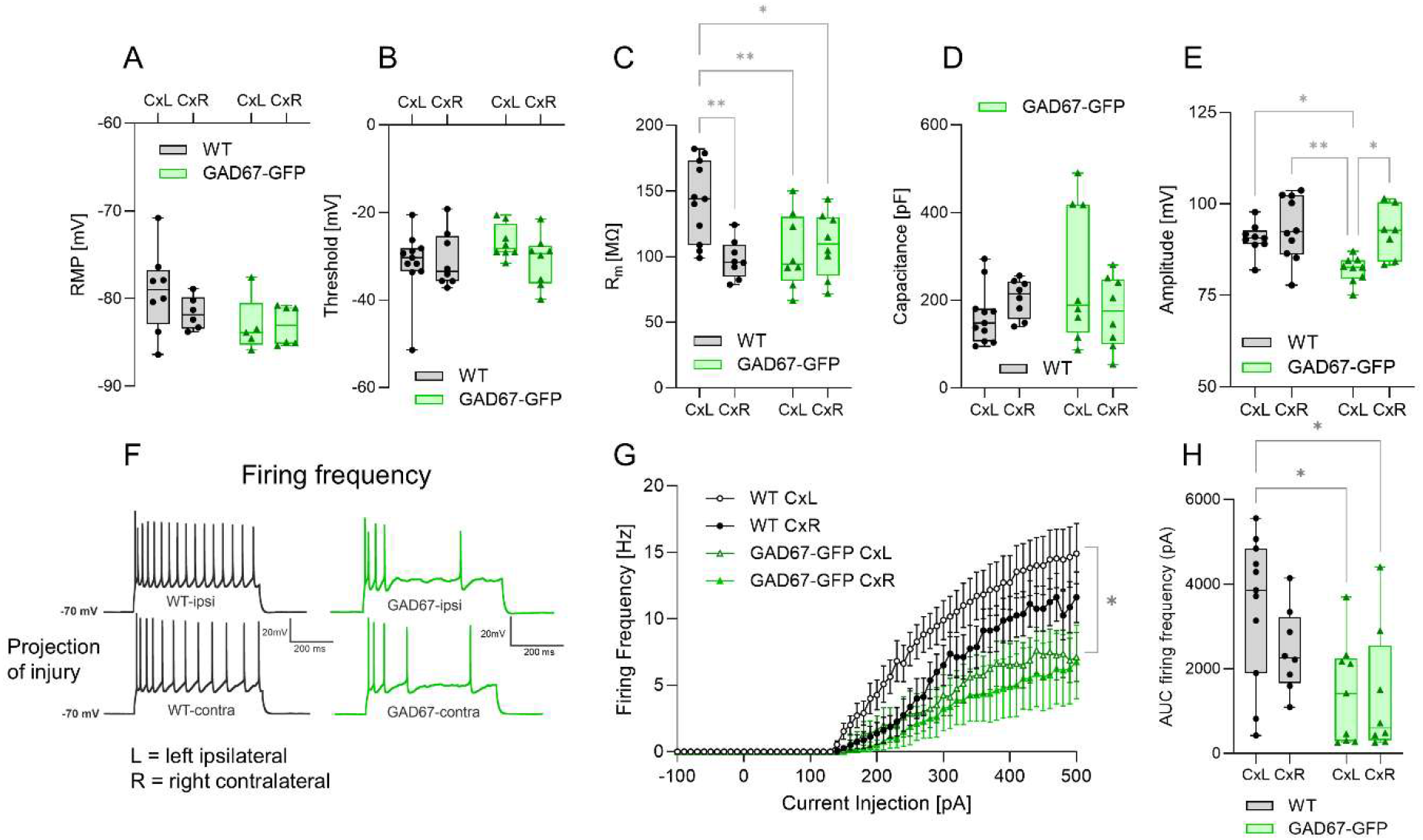
Whole cell patch-clamp recordings of pyramidal neurons in the left (ipsilateral, non-projection) and right contralateral cortical hemisphere in GAD67-GFP and wildtype mice (n = 3/3 female and male for each genotype) in acute cortical slices 10-14 days after SNI. The left somatosensory cortex is ipsilateral to the injured nerve, but nociceptive input crosses to the opposite side in the spinal cord. Hence, the right contralateral cortex receives the direct input from the injured left sciatic nerve. **A:** Unchanged resting membrane potential (RMP); **B:** Unaltered action potential threshold **C, D:** Membrane resistance (R_m_) and capacitance (C_m_); **E:** Action potential amplitudes at rheobase; Note the decreased membrane resistance in the right contralateral hemisphere (projection cortex) after SNI as compared to the non-projection left ipsilateral cortex in wildtype mice. In GAD67-GFP mice, membrane resistances in both hemispheres resembled the low right-sided post-SNI resistance of the wildtype mice suggesting a partial decrease in the intrinsic excitability, and hence failure to adapt to SNI. **F, G, H:** Representative voltage traces and mean AP firing frequency of pyramidal neurons upon current injection, and areas under the curves (AUC) for statistical comparison. The firing frequency was significantly reduced in GAD67-GFP cortices in both hemispheres. Data were compared with two-way ANOVA for the within subject factor “current” or “side” and the between subject factor “genotype” and subsequent posthoc analysis using an adjustment of alpha according to Tukey. The boxes show the interquartile range, the line is the median, the whiskers show minimum to maximum and the scatters are individual neurons of 6 mice per genotype. Asterisks indicate statistically significant differences. *P<0.05; **P<0.01.

Analyses of spontaneous excitatory and inhibitory postsynaptic currents (sEPSCs and sIPSCs) support this conclusion (Figure 6). In WT slices, a trend for a reduction in the sEPSC frequency could be observed (p=0.0562), pointing at a partial loss of synaptic input to this side due to cut of the tibial nerve. In contrast, the sIPSC frequency remained unaltered in both genotypes after injury also compensating the mild reduction in the sEPSC frequency in WT slices, as indicated by an unaltered E/I ratio of synaptic input frequencies (Figure 6). The data suggest that partial loss of peripheral input in WT shifted excitatory synaptic transmission toward fewer sEPSCs, which, however, did not affect the overall E/I ratio within the network. In GAD67-GFP mice, the sEPSC frequency did not change after injury, suggesting that the GAD67-GFP occupies a pre-adapted, ‘post-injury–like’ synaptic state and fails to engage the normal deafferentation-induced synaptic or interhemispheric remodeling.

**Figure 6.**
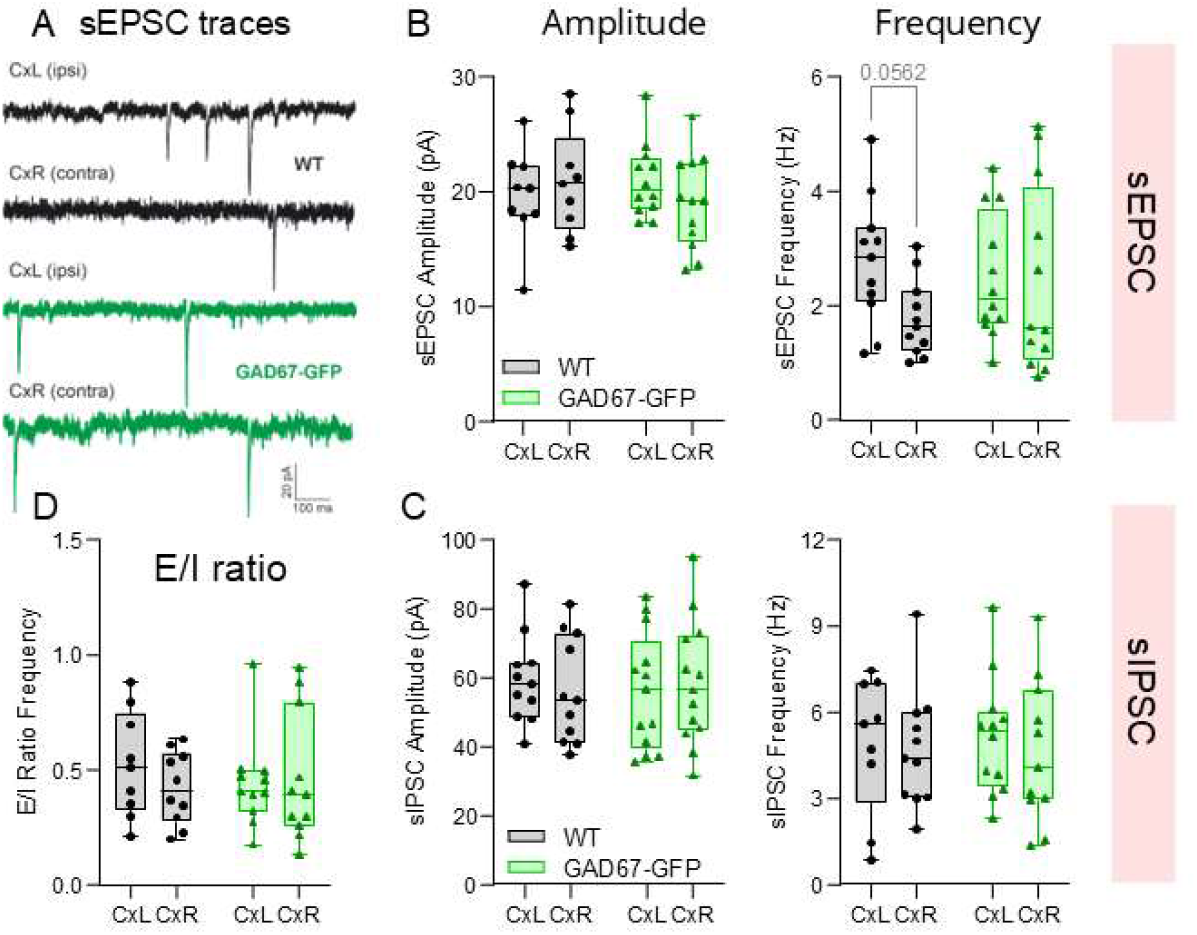
Spontaneous excitatory and inhibitory postsynaptic currents (sEPSC, sIPSC) of pyramidal neurons in ex vivo cortical slices. The slices were prepared as described in figure 5. The upper row shows sEPSC features, the bottom row the corresponding sIPSC features and E/I ratios for frequency. sEPSC and sIPSC signals were recorded for at least 5 min, and for each neuron, ≥150 events were included for amplitude and kinetic analysis to obtain the cell’s average. **A:** sEPSC traces, **B:** sEPSC amplitude and frequency, **C:** sIPSC amplitude and frequency, **D:** E/I frequency ratio. Data were submitted to two-way ANOVA for the within subject factor “side” and the between subject factor “genotype” and subsequent mutual posthoc analysis using an adjustment of alpha according to Tukey. The boxes show the interquartile range, the line is the median, the whiskers show minimum to maximum and the scatters are individual neurons of 6 mice per genotype. Comparisons with P<0.1 show the P-value.

Our previous study on juvenile GAD67-GFP cortical slices revealed a paradoxical tonic GABAergic inhibition despite GABA deficiency [48]. We further found in that study an increased ambient GABA via GAT3 mediated reverse outward transport. Although not experimentally re-tested, we assume that this reverse GAT3 transport was still operative in adolescent mice providing an explanation for the relative mild behavioral phenotype that manifested only after nerve injury with a subtle nociceptive hypersensitivity but otherwise completely normal behavior.

In support of this conclusion, cortical network recordings of synaptically-evoked field potentials (FPs) in acute slices of GAD67-GFP mice in our MEA-setup showed larger amplitudes of evoked FPs in the ipsilateral left hemisphere than in the contralateral right projection hemisphere, where field potentials were reduced (Figure 7). In wildtype mice, SNI did not evoke such opposing regulations of FPs in ipsi - versus contralateral hemispheres. Indeed, ipsi- and contralateral field potentials were almost equal in wildtype slices. The strong rise of amplitudes in the ipsilateral hemisphere in GAD67-GFP slices i.e. in the hemisphere not receiving direct input from the injured side might represent a compensatory mechanism reminiscent of cortical diaschisis following traumatic brain injury [37].

**Figure 7.**
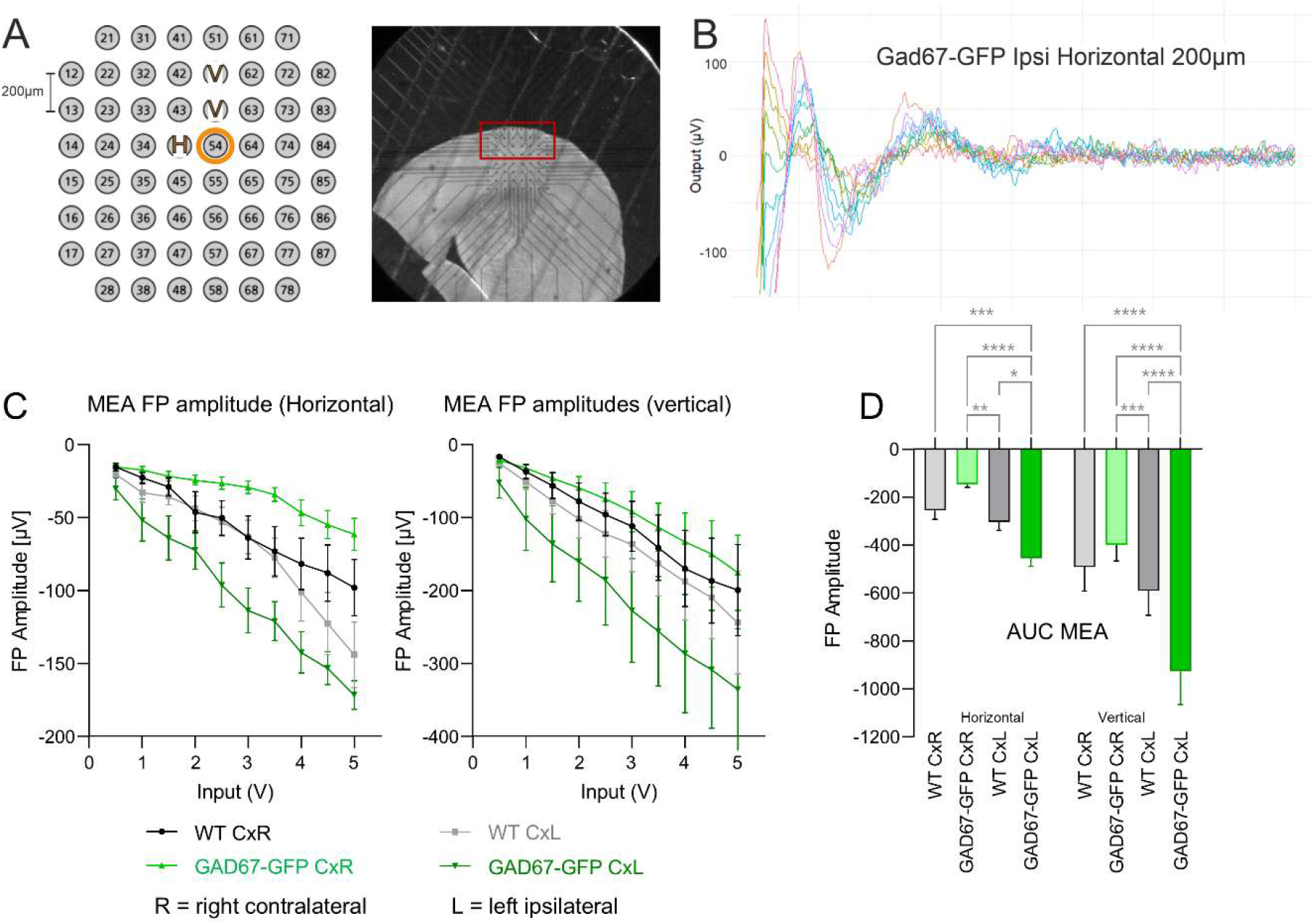
Cortical network activity in GAD67-GFP mice in MEA-recordings. Extracellular field potentials were recorded using multielectrode arrays (MEA) on freshly prepared slices of GAD67-GFP and wildtype control mice (n = 3/3 female and male for each genotype) 10-14 days after left-sided sciatic nerve injury (SNI). **A, B:** Illustration of the MEA chip on top of the brain slice showing the positioning of the electrodes in the area of the somatosensory cortex (SSC) and exemplary traces upon horizontal stimulation. The representative recording was obtained from electrode 54, with stimulation from a neighboring electrode with a distance of 200 µm either in horizontal (H) or vertical (V) direction. Only the 200 µm distant stimulation was used for analysis. **C:** MEA field potential (FP) amplitudes at increasing voltage stimulations for horizontal and vertical stimulation directions. The data represent means and sem from recordings of both ipsilateral left and contralateral right hemispheres (projection side) of GAD67-GFP and wildtype mice. For statistical analyses of the FP versus input curves, areas under the curve (AUC) were calculated using the linear trapezoidal rule. **D:** Area of the FP-amplitudes recorded from the two different inputs (AUC) were analyzed with one-way ANOVA separately for horizontal and vertical stimulation, and subsequent posthoc analysis using a correction of alpha according to Šidák. Asterisks indicate statistically significant differences between groups, *P<0.05, **P<0.01, ***P<0.001, ****P<0.0001.

### Metabolic adaptations to GABA deficiency and peripheral nerve injury

Metabolomic studies of the ipsilateral and contralateral cortex and lumbar spinal cord confirmed the expected modest deficiency of GABA in GAD67-GFP mice (Figure 8-9). In wildtype mice, but not in GAD67-GFP mice, GABA was increased in the ipsilateral spinal cord receiving direct input from the injured nerve as compared to the contralateral side (Figure 8A). Detailed analyses of the metabolome showed that low levels of GABA were accompanied with low levels of carnosine (beta-alanyl-L-histidine) (Figure 8B), cystathionine (8C), glutathione (8D), tryptophane and glucose-6-phosphate (Figure 9), which are all interconnected in carnosine and GABA biosynthesis pathways (Figure 10). Carnosine is a naturally occurring dipeptide with high concentrations in muscle and the brain. It is advertised as a dietary supplement with antioxidant, anti-inflammatory, and pH buffering effects, and is supposed to protect against oxidative stress, generation of glycation end products, and muscle fatigue [62, 63]. However, rigorous efficacy studies are lacking, and oral supplementation is unlikely to affect CNS levels. Cystathionine is a crucial intermediate amino acid in the trans-sulfuration pathway, formed from homocysteine and serine, primarily serving as the precursor to cysteine, a building block for glutathione, which was also reduced. It is vital for methionine metabolism and cellular redox balance. Via glutathione and hydrogen sulfide (H_2_S), cystathionine is supposed to maintain cellular protection against oxidative stress [64, 65].

**Figure 8.**
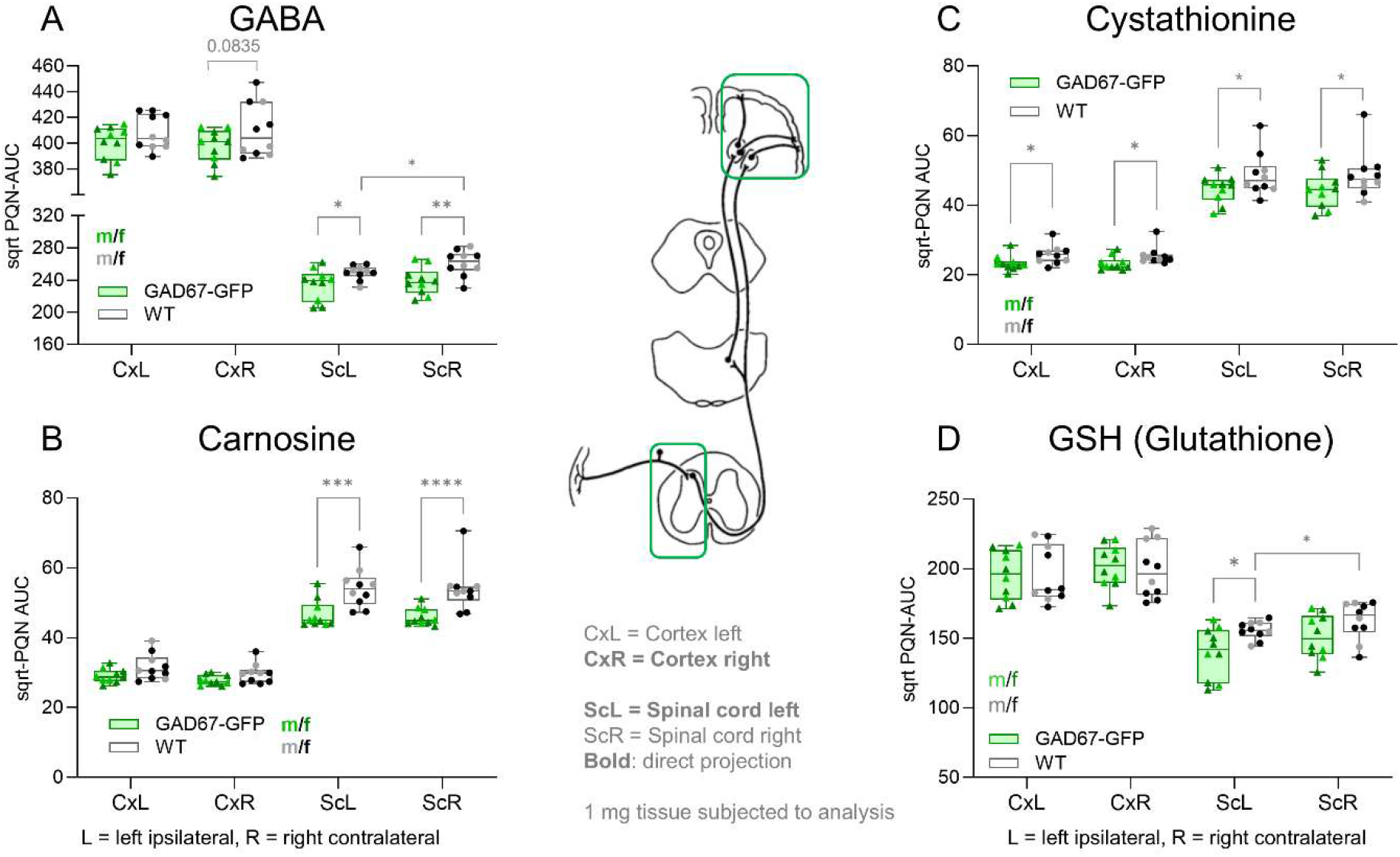
**A, B:** Box/scatter plots of GABA and carnosine normalized AUCs of UHPLC-mass spectrometry analyses in 1 mg of right (contralateral) and left (ipsilateral) cortices and right and left lumbar spinal cord. AUCs were normalized according to the Probabilistic Quotient Normalization (PQN) method to adjust for tissue-related differences (cortex, spinal cord) and square root transformed (sqrt) to linearize the areas. The boxes show the interquartile range, the line is the median, whiskers show minimum to maximum, the scatters are the mice. Please note that the lighter scatters are the males and the darker scatters are females (n = 5/5 male/female for each genotype). Data were compared with 2-way ANOVA followed by posthoc analysis for side (left, right) and genotype. Asterisks show statistically significant between-group differences, *P<0.05, **P<0.01, ***P<0.001, ****P<0.0001. **C, D:** In analogy to A, B the graphs show box/scatter plots of normalized AUCs of two antioxidative metabolites, cystathionine and glutathione (GSH) which were significantly downregulated in GAD67-GFP mice as compared to wildtype. The boxes show the interquartile range, the line is the median, the whiskers show minimum to maximum. Statistics as in A, B.

**Figure 9.**
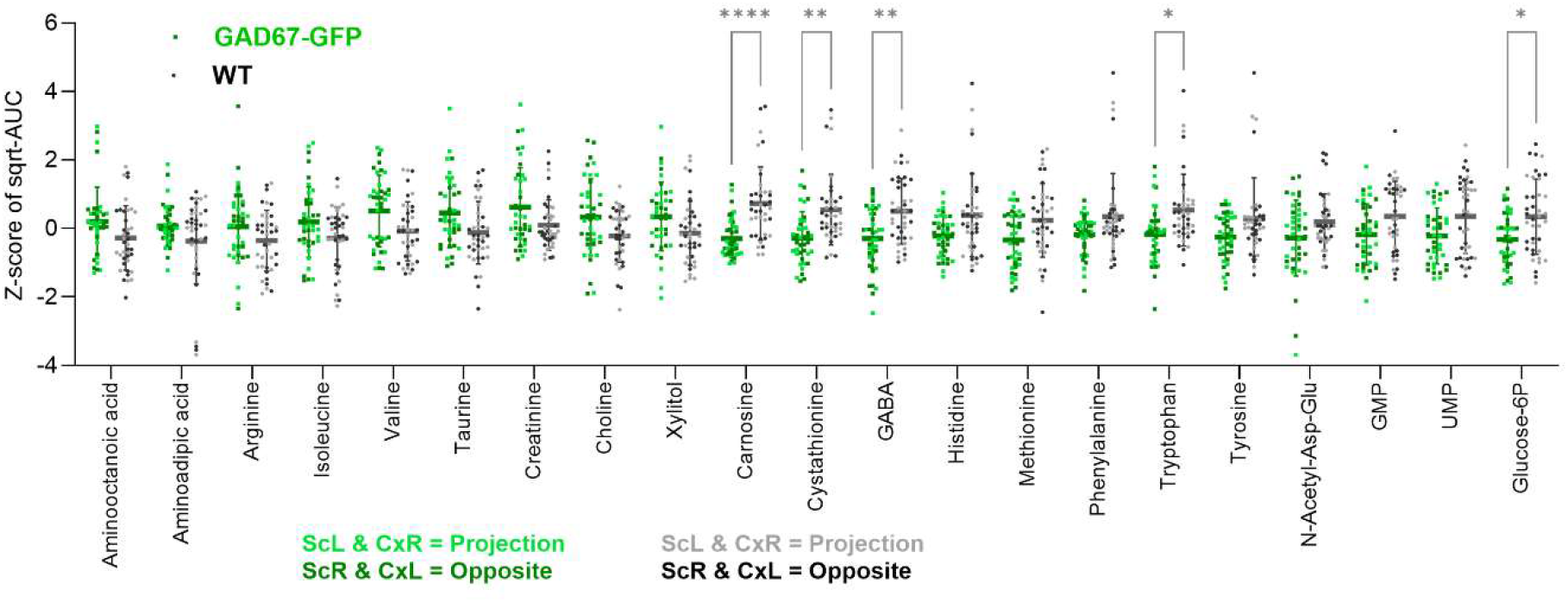
Overview of candidates metabolites which were obtained from UHLPC-HRMS metabolome analyses in right (contralateral) and left (ipsilateral) cortices and right and left lumbar spinal cords of n = 10 mice per genotype. To reveal even subtle genotype dependent differences, metabolites from different sites were pooled and transformed to z-scores (autoscaling) to have a common median and variance of 1. The sites are color-coded, light and dark green for GAD67-GFP; grey and black for WT. Data were compared with 2-way ANOVA and subsequent posthoc analysis according to Šidák for genotype. The scatters show the Z-transformed AUC data per mouse and site, i.e. each mouse is represented by 4 scatters. Asterisks show statistically significant between-group differences, *P<0.05, **P<0.01, ****P<0.0001

**Figure 10.**
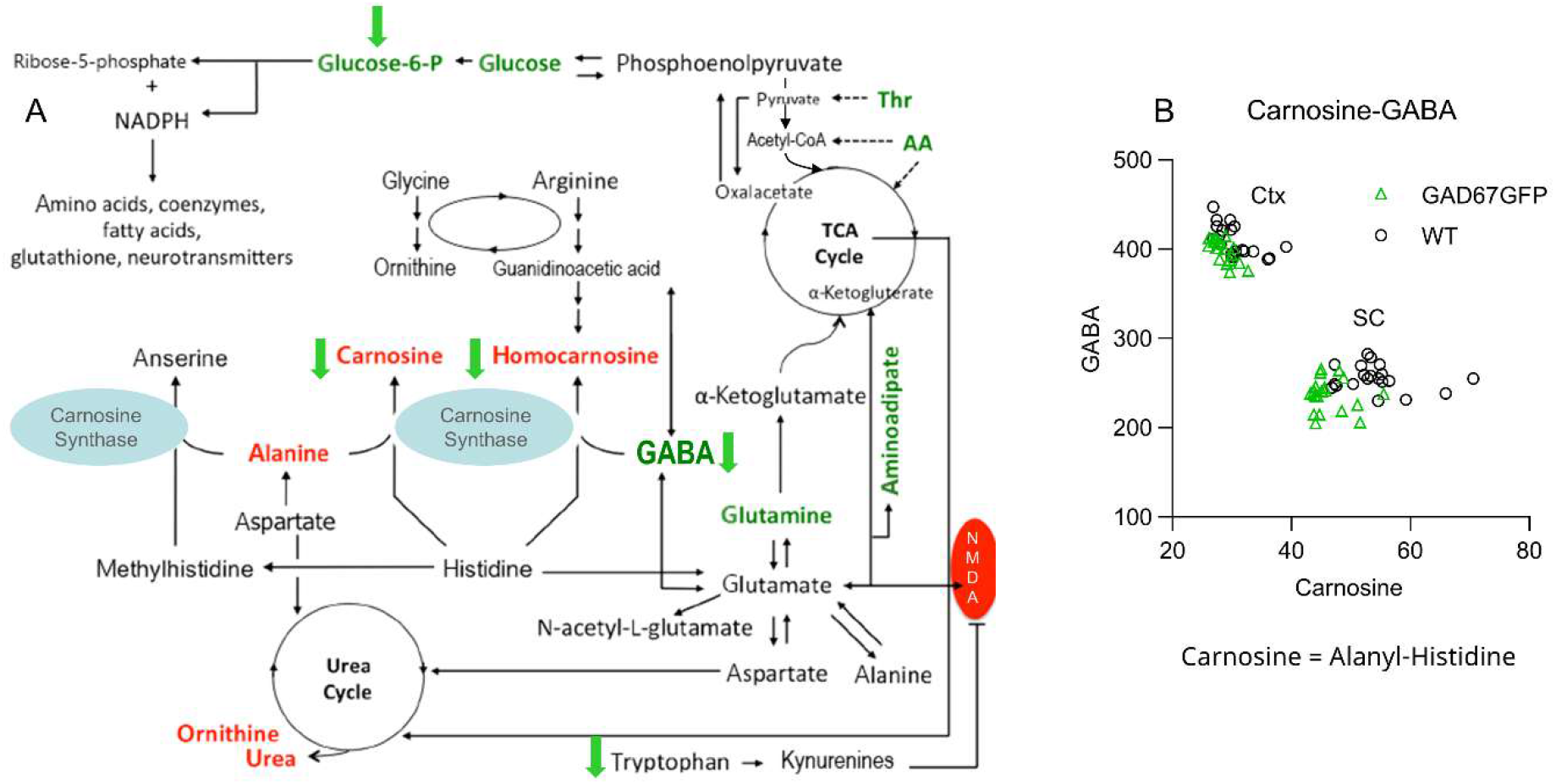
Illustration of the metabolic pathways involved in GABA homeostasis **A:** The metabolic pathways or candidates which were regulated in GAD67-GFP mice are highlighted with a green arrow indicating up or downregulation in association with or in consequence of GABA deficiency (adapted from [88]). **B:** Carnosine versus GABA scatter plots show that the genotypes can be differentiated by these two metabolites. The differences are stronger in the spinal cord.

Lipidomic analyses showed distinct lipid profiles for spinal cord and brain, particularly for fatty acids, sphingolipids and lyso-species, but did not reveal any differences between GAD67-GFP and WT mice, neither in the lumbar spinal cord nor cortex (Suppl. Figure S1, source data available at Biostudies with the accession number S-BSST2724). The comparison shows that the mild reduction of GABA synthesis has a mild metabolic impact specifically on GABA pathways but does not cause critical changes of lipid or polar metabolic building blocks or unrelated metabolic pathways. The equality of lipidomic patterns of ipsilateral and contralateral spinal cord also shows that the peripheral injury did not cause myelin degradation or back-dying of neurons. We therefore infer that the observed differences in post-SNI thermal nociceptive hypersensitivity are caused by differences of GABA itself, possibly contributed by reduced antioxidative capacity owing to low carnosine, cystathionine and glutathione.

## Discussion

We show that haploinsufficiency of glutamate decarboxylase (GAD67) induces a GABA deficit in the spinal cord and weaker in the cortex of adolescent mice, which is counterbalanced by adaptive adjustments in GABA metabolism and transport to preserve synaptic E/I homeostasis. While our data suggest that the adaptation was successful in preventing hyperexcitability in naïve uninjured healthy mice, this compensatory capacity appears exhausted when the system is challenged by sciatic nerve injury resulting in thermal nociceptive hypersensitivity and allodynia in GAD67-GFP mice that was not detected in wildtype control mice in the unbiased TGR maze-test that measures thermal place preference behavior rather than mechanically or heat stimulated paw withdrawal. The latter, more commonly used nociceptive assays rely on manually scoring paw-withdrawal latencies upon plantar stimulation, introducing observer bias and requiring confinement of the mouse in a small grid-floor enclosure associated with fear and stress that may occasionally elicit un-stimulated paw withdrawal by positioning the stimulator underneath the paw. Hence, our results neither contradict findings from stimulated paw-withdrawal assays nor imply that the tibial-only SNI was unsuccessful. Correct placement of the proximal ligature and integrity of the nerve stump were verified by necropsy and histological examination of the sciatic nerve. Nevertheless, one-branch SNI may induce weaker or less consistent hypersensitivity, resulting in greater interindividual variability, although previous studies report that single-branch SNI is as effective as the standard neuropathic pain model [53, 54]. In our cohort, five wildtype mice showed a transient loss of sensitivity rather than hypersensitivity, such that mean preference temperatures remained unchanged. Sensory loss is known to occur during the first days after SNI and is typically no longer detectable by paw-withdrawal testing at 7 dpi, although it may persist in some animals. Importantly, none of the GAD67-GFP mice displayed behavior consistent with such “pain-protective” numbness.

In GAD67-GFP hemizygous mice, chronic reduction of GABA synthesis appears to be counterbalanced by a tonic inhibitory influence on cortical pyramidal neurons, as suggested previously in brain slices of juvenile mice [48] and now evident as a reduced firing capacity in adolescent female and male mice. This tonic brake may arise from enhanced ambient GABA signaling via transporter-dependent non-vesicular release [66, 67] and/or homeostatic upregulation of intrinsic conductance such as hyperpolarization-activated cation currents, Ih [68, 69], which together limit pyramidal excitability at baseline and may protect against spontaneous nociceptive hypersensitivity [70, 71]. However, because inhibition is already tonically engaged, the system appears to have little dynamic range left to further increase inhibitory control when peripheral input becomes abnormal after sciatic nerve injury. As a consequence, nerve injury may lead to a relative failure of activity-dependent inhibitory upscaling and manifesting as network hyperactivity [72-74], which was most evident in the non-projection (ipsilateral) cortex where sensory drive was preserved, whereas the projection cortex itself showed reduced activity possibly due to loss of afferent input. Hence, GAD67 deficiency during adolescence plus nerve injury did not simply increase cortical excitability but rather created an interhemispheric and microcircuit-level imbalance that may favor maladaptive pain processing.

In short, GAD67 haploinsufficiency paradoxically appears to lock cortical inhibition into a tonic mode that stabilizes cortical excitability at rest but leaves little capacity for further inhibitory recruitment after nerve injury. When peripheral input becomes abnormal, inhibition cannot scale up, leading to hyperactive networks in the intact hemisphere while the deafferented projection cortex remains more quiet. For development or persistence of “pain” it is likely not (only) net activity in the projection-side SSC but rather a shift in interhemispheric balance that determines the outcome. A putative scenario could be that the non-projection ipsilateral side becomes relatively more active, because of the partial denervation of the projection side, and a recruitment of normally silent circuits [75]. The resulting imbalance could distort bilateral integration of nociceptive and somatosensory information [76] and favor maladaptive representations of the injured limb. If the projection-side cortex is relatively suppressed, it may fail to exert normal top-down or lateral inhibitory control over subcortical nociceptive circuits, while the other side becomes hyper-responsive to aberrant inputs e.g., from callosal or higher-order thalamic pathways [77]. In addition, microcircuit or temporal inhibition may be redistributed between interneuron subtypes, although summary E/I ratios were unchanged.

Our previous work in traumatic brain injury models demonstrated that interhemispheric balance is a critical determinant of successful cortical adaptation. For this reason, we focused here on comparing the two hemispheres within the same animals rather than comparing SNI and sham groups. In the spinal cord, ipsilateral and contralateral inputs are clearly segregated, but the cortex receives convergent information from both sides of the body. Although the major ascending pathways cross and predominantly project to the contralateral cortex, a minor subset of ipsilateral projections, such as components of the spinothalamic tract, also reaches the cortex [78, 79]. Consequently, the hemisphere ipsilateral to the peripheral nerve injury may not represent a completely “normal” reference. This constitutes a limitation of our study, as we did not include an additional sham-operated control group. For the cortical metabolomic analyses, we used historical mice of comparable age and sex as reference controls and did not detect metabolic differences between the ipsilateral cortex of WT SNI mice and the corresponding tissue from historical naïve animals (data not shown). Thus, at the metabolic level, the ipsilateral WT cortex can be considered an appropriate “normal” reference.

A further layer of vulnerability in GAD67-GFP mice emerges from the metabolomic profile of cortex and spinal cord. Beyond the partial GABA deficit which was obvious in the spinal cord but not significant between cortices at a “side by temporal” resolution, haploinsufficiency was accompanied by a coordinated reduction in several metabolites with established roles in redox buffering and cellular stress resistance, including carnosine, cystathionine, and glutathione [63]. These changes occurred bilaterally and were present already at baseline, indicating that the metabolic milieu of GAD67-deficient tissue is shifted toward lower antioxidant capacity even before injury. It is therefore not possible to distinguish between the primary effects of GABA deficiency and the secondary effects of the established compensations, and the response to nerve injury is not a direct reflection of “GABA deficiency” in isolation, but rather a reaction from a “rewired” nervous system.

In wildtype mice, sciatic nerve injury triggers an ipsilateral increase in GABA levels within the spinal cord [55], a response thought to support adaptive inhibitory reinforcement and to limit the transition to chronic pain. In contrast, we now show that GAD67-GFP mice fail to mount this injury-induced GABA upregulation, suggesting that the system is already operating near its ceiling and cannot further boost inhibitory tone when challenged. The concomitant depletion of carnosine, cystathionine, and glutathione may exacerbate this limitation by reducing the ability of neurons and glia to buffer oxidative stress generated by peripheral nerve injury [51, 80, 81] and heightened network activity [82, 83]. Oxidative stress is known to impair GABAergic signaling at multiple levels—from interneuron excitability to vesicular release and receptor function [84-86]—and thus a reduced redox reserve could further constrain inhibitory plasticity precisely when it is most needed. Together, the metabolic and neurotransmitter deficits may create a state in which both the biochemical substrates for inhibition and the cellular resilience mechanisms that normally support adaptive responses to injury are compromised.

## Supporting information

Supplemental Figure with legends

Supplemental Analytical Procedures

TGR video GAD67-GFP after SNI

TGR video GAD67-GFP before SNI

TGR video WT after SNI

TGR video WT before SNI

## Statements and declarations

### Animal Ethics Approval

The experiments were approved by the local Ethics Committee for Animal Research (Darmstadt, Germany) (V54 19c 20/15 FK1110) and the Landesuntersuchungsamt Rheinland-Pfalz (for in vitro electrophysiology), and they adhered to the European guidelines and to those of GV-SOLAS for animal welfare in science and agreed with the ARRIVE guidelines.

### Consent for publication

All authors have approved the manuscript for publication.

### Data availability statement

Lipidomic and metabolomic processed data were deposited at Biostudies [87] with the accession number S-BSST2724. The data is still private but can be accessed via the link: https://www.ebi.ac.uk/biostudies/studies/S-BSST2724?key=4b50c546-7752-4a09-9cbf-5cba96b61d1c

### Author contributions

JS did behavioral and immunofluorescence studies, CH and CAS did electrophysiology studies, MPW and LF supervised behavioral studies and contributed to tissue sample collection, LH and RG supervised and analyzed metabolomics analyses, TM and IT conceived the experiments, acquired funding and Ethical Approval, and supervised studies. IT did the surgery, analyzed behavioral and metabolomic data, made the figures and drafted the manuscript. All authors contributed to editing the manuscript and approved submission of the latest version.

## Acknowledgements

We thank Carlo Angioni (Institute of Clinical Pharmacology, Goethe-University, Frankfurt, Germany) and Rahmat Mojaradfar (Fraunhofer ITMP, Frankfurt, Germany) for assistance in UPLC-HRMS measurements.

## Supplementary videos

The videos show 1-2 min recordings of mouse behavior in the first quarter of the Thermal Gradient Ring (TGR) test before and after SNI in wildtype and in GAD67-GFP mice. File names indicate the genotype and time point.

