## Supplemental Figure with legends for "Nerve injury triggers nociceptive hypersensitivity with interhemispheric divergence in haplodeficient GAD67-GFP mice"

Suppl. Figure S1

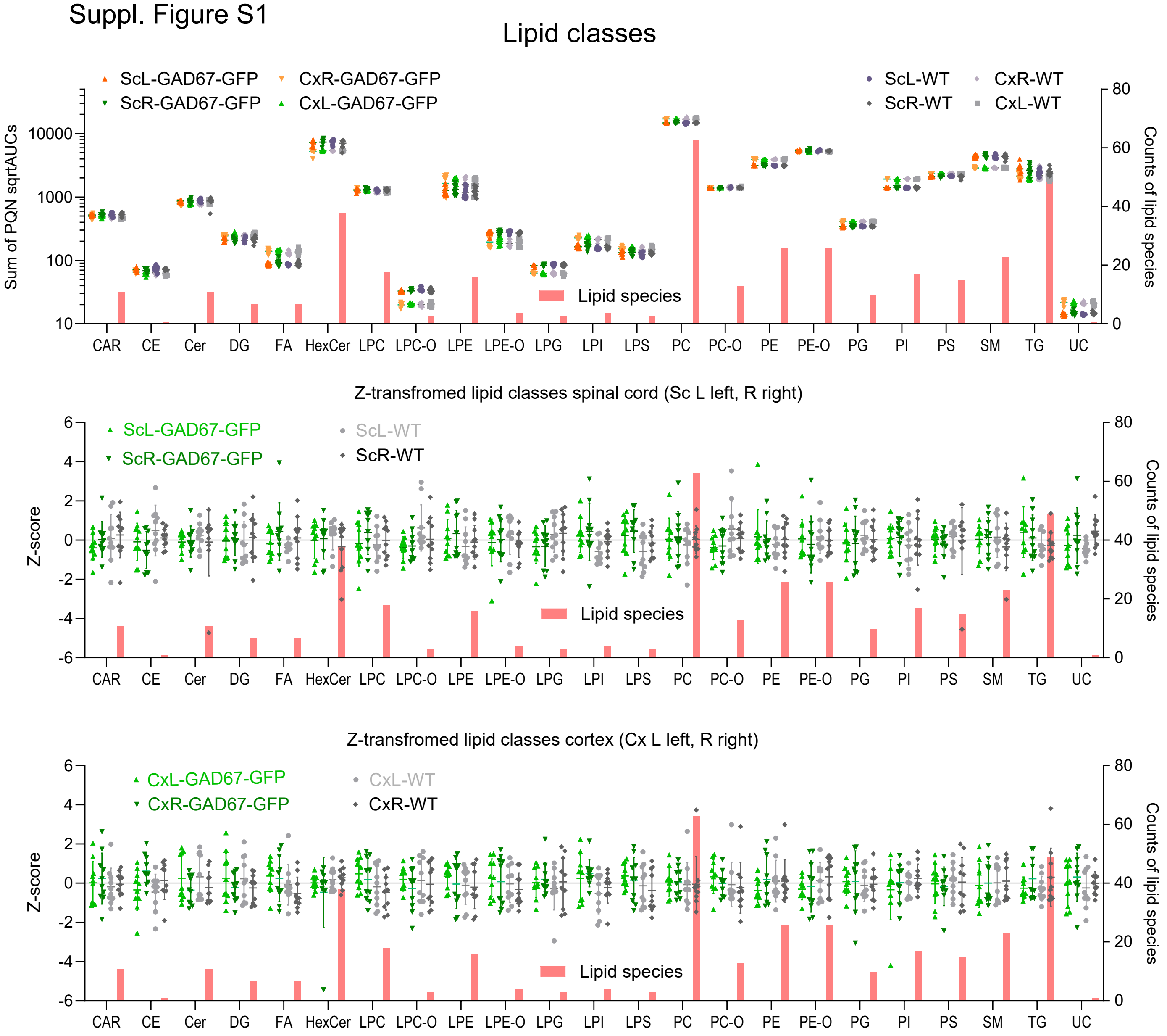


Suppl. Figure S1

Overview of lipid classes obtained from UHLPC-MS/MS lipidome analyses in right (contralateral) and left (ipsilateral) cortices and right and left lumbar spinal cords of n = 10 mice per genotype (each 5/5 male/female). Tissue was obtained 14 days after injury of the **left** tibial nerve. The top panel shows the sums of sqrt AUCs from lipid species of the respective class. The counts of lipid species per class are scaled on the right Y-axis. To reveal between side and between genotype differences, lipid classes from spinal cord and cortex were transformed to z-scores (autoscaling) to have a common mean and variance of 1. Sites and genotypes are color-coded, light for cortex and darker for spinal cord, yellow/green for GAD67-GFP and grey/black for WT. The scatters show the summed AUCs or Z-transformed AUCs per mouse and site. The line is the average, the whiskers are SD. Spinal cord and cortex have distinct lipid profiles (top panel), but there was no statistically significant difference between ipsilateral and contralateral sides or between GAD67-GFP and wildtype mice.

Abbreviations: CAR, acylcarnitines; Cer, ceramides; CE, cholesterol ester; DG, diglycerides; FA, fatty acids; HexCer, hexosylceramides; LPC, lysophosphatidylcholines; LPE, lysophosphatidylethanolamines; LPG, lysophosphatidylglycerols; LPI, lysophosphatidylinositols; LPS, lysophosphatidylserines; PC, phosphatidylcholines; PE, phosphatidylethanolamines; PG, phosphatidylglycerols; PI, phosphatidylinositols; PS, phosphatidylserines; SM, sphingomyelins; TG, triglycerides; –O ether bound lipids
